# Simulation-Based Inference of Cell Migration Dynamics in Complex Spatial Environments

**DOI:** 10.1101/2025.09.25.678460

**Authors:** Jonas Arruda, Emad Alamoudi, Robert Mueller, Marc Vaisband, Ronja Molkenbur, Jack Merrin, Eva Kiermaier, Jan Hasenauer

**Affiliations:** Life & Medical Sciences Institute, University of Bonn, Bonn, Germany; Bonn Center for Mathematical Life Sciences, University of Bonn, Bonn, Germany; Center for Interdisciplinary Digital Sciences, Department Information Services and High Performance Computing, TUD Dresden University of Technology, Dresden, Germany; Life & Medical Sciences Institute, Immune and Tumor Biology, University of Bonn, Bonn, Germany; Institute of Science and Technology Austria, Klosterneuburg, Austria; Department of Medicine 1/CITABLE, Friedrich-Alexander University Erlangen-Nürnberg, Erlangen, Germany; Deutsches Zentrum Immuntherapie (DZI), Universitätsklinikum Erlangen, Erlangen, Germany

## Abstract

To assess cell migration in complex spatial environments, microfabricated chips, such as mazes and pillar forests, are routinely used to impose spatial and mechanical constraints, and cell trajectories are followed within these structures by advanced imaging techniques. In systems mechanobiology, computational models serve as essential tools to uncover how physical geometry influences intracellular dynamics; however, decoding such complex behaviors requires advanced inference techniques. Here, we integrated experimental observations of dendritic cell migration in a geometrically constrained microenvironment into a Cellular Potts model. We demonstrated that these spatial constraints modulate the motility dynamics, including speed and directional changes. We show that classical summary statistics, such as mean squared displacement and turning angle distributions, can resolve key mechanistic features but fail to extract richer spatiotemporal patterns, limiting accurate parameter inference. To solve this, we applied neural posterior estimation with in-the-loop learning of summary features. This learned summary representation of the data enables robust and flexible parameter inference, providing a data-driven framework for model calibration and advancing quantitative analysis of cell migration in structured microenvironments.

## Introduction

Tissues are complex, highly organized systems comprising cells and extracellular components that exhibit dynamic behavior and structural heterogeneity across developmental stages. This spatiotemporal complexity, encompassing both biochemical gradients and mechanical cues, critically influences cell behavior, particularly migration dynamics within structured or constrained environments [1, 2]. Dendritic cells (DCs) belong to the group of innate immune cells, which activate T cells via antigen presentation [3]. To efficiently reach their final destination within the secondary lymphoid organs, DCs follow chemical guidance cues presented as chemokine gradients across the lymphatic vasculature [4]. Directional cell migration in response to chemotactic cues is a cornerstone of many (patho)-physiological processes, including immune surveillance and response [5], inflammation [6], and cancer metastasis [7, 8, 9]. This guided movement, termed chemotaxis in response to soluble attractants or haptotaxis when driven by immobilized chemokine gradients, involves the binding of chemokines to receptors on the cell surface, which activates intracellular signaling pathways that coordinate directed migration by integrating chemotactic sensitivity, migration persistence, cell morphological dynamics, and the traction forces modulated by the mechanical and chemical microenvironment [10]. Advances in live cell imaging have facilitated the acquisition of extensive quantitative data to capture the dynamic behavior of migrating cells [11]. The interpretation of these data to elucidate pattern formation and tissue dynamics necessitates advanced computational modeling due to the complexity of the data. Accordingly, computational models have become valuable tools for capturing the multiscale interactions and emergent behaviors that characterize biological systems. Modeling approaches, including discrete and continuous models in both temporal and spatial domains, as well as hybrid frameworks, have been employed to capture cell behavior and cell-cell interactions within complex environments [12, 13, 14, 15, 16]. Specifically, Cellular Potts models [17] and vertex models, which are continuous in time and discrete in space, are extensively utilized to simulate cell interactions, with numerous software platforms available for implementing and testing these models [18, 19, 20, 21, 22, 23]. Various studies have leveraged simulations to unravel the dynamics of chemotaxis and cell migration, emphasizing the pivotal role of computational models in understanding cellular behavior [24, 25, 26].

A central challenge in applying these models is their calibration using experimental data. Typically, model parameters are not directly measurable, yet they are crucial for elucidating biological mechanisms. Parameter estimation enables quantitative modeling [27, 28], hypothesis testing [29], and prediction of dynamic responses to perturbations [30, 31]. The application of parameter estimation methods to cell migration processes has been previously explored to infer parameters governing cell motility and intercellular forces [32, 33]. In particular, parameter inference methods, such as approximate Bayesian computation (ABC), facilitate model calibration based on simulations without the need to derive a likelihood, which can be prohibitively complex for Cellular Potts models [34, 35, 36].

State-of-the-art ABC pipelines typically rely on summary statistics to reduce the dimensionality of the data. These statistics, such as the mean squared displacement or turning angle distributions of cell movement, are often hand-crafted based on domain knowledge. However, they may fail to provide relevant information for precise parameter estimation. To address these challenges in the context of ABC, recent approaches have included adaptive algorithms that weight hand-crafted summaries to maximize the posterior information gained from a dataset or to learn summaries on pairs of parameters and simulations [37, 38, 39]. However, handcrafted summaries or summaries based on the posterior mean may not be sufficient to reconstruct the posterior shape [40, 39].

Alternatives to ABC, such as neural posterior estimation (NPE) methods, employ neural networks to learn mappings from data to model parameters, effectively handling high-dimensional data and sophisticated models and circumventing the limitations of conventional ABC methods [41]. By training neural networks, relevant features can be extracted automatically from the data, neural posterior estimation can improve parameter estimation accuracy [42].

In this study, we aimed to improve the inference of spatial cell movement models in complex environments by applying both classical and machine learning simulation-based inference methods. A comprehensive inference pipeline was implemented to evaluate the different inference approaches. As an important example of the interplay between spatial structure and cell migration, we focused on dendritic cell movement in complex environments. Therefore, we designed microstructured polydimethylsiloxane chips that impose obstacles on migrating dendritic cells and recorded their trajectory data. To analyze the data, a Cellular Potts model of migration in a spatially complex environment was constructed, and approximate Bayesian computation with different summary statistics was compared with neural posterior estimation using simulated and experimental data. We demonstrated the ability of the model to capture the modulation of speed and direction through obstacles and chemokine gradients; hence, we present a data-driven framework to analyze how environmental mechanics and structure influence cell behavior, thereby advancing computational approaches in systems mechanobiology.

## Results

To quantitatively dissect how chemokine gradients and physical constraints influence dendritic cell migration, we designed a structured *in vitro* assay and built a mechanistic model *in silico* based on the Cellular Potts framework. Using synthetic data with known ground truth, we systematically evaluated four simulation-based inference strategies: (i) approximate Bayesian computation (ABC) with hand-crafted summary statistics (displacement, velocity, turning angle, and angle degree); (ii) ABC employing a neural network as a feature extractor trained either to predict the posterior mean (ABC-PM) or (iii) with summaries tailored for inference (ABC-NPE); and (iv) neural posterior estimation (NPE) with jointly trained summary networks (as described in the Methods). After identifying NPE as the most flexible approach, we applied this strategy to calibrate our model on experimentally tracked dendritic cell trajectories under chemotactic stimulation within a microstructured polydimethylsiloxane (PDMS) pillar array. The resulting model faithfully reproduced the observed migration patterns and enabled realistic simulations under varying environmental conditions, offering mechanistic insights into how chemical signals and spatial barriers jointly govern immune cell navigation.

### Dendritic cell migration in pillar forests

Dendritic cells must efficiently migrate through complex tissue environments to reach lymphoid organs, guided by chemokine gradients. Understanding how they integrate chemical signals with physical constraints is critical for deciphering the mechanisms of the immune system.

To study this interplay, we exposed bone marrow–derived dendritic cells to a stable CCL19 gradient and confined them within a microfluidic chip containing a pillar forest with 10 µm gaps (Figure 1A and [43]). Time-lapse videomicroscopy was used to track the nuclear positions at 30-second intervals in the visible window (Figure 1B), yielding trajectories of variable length depending on how long a cell remained within the field of view (further experimental details in Methods).

**Figure 1:**
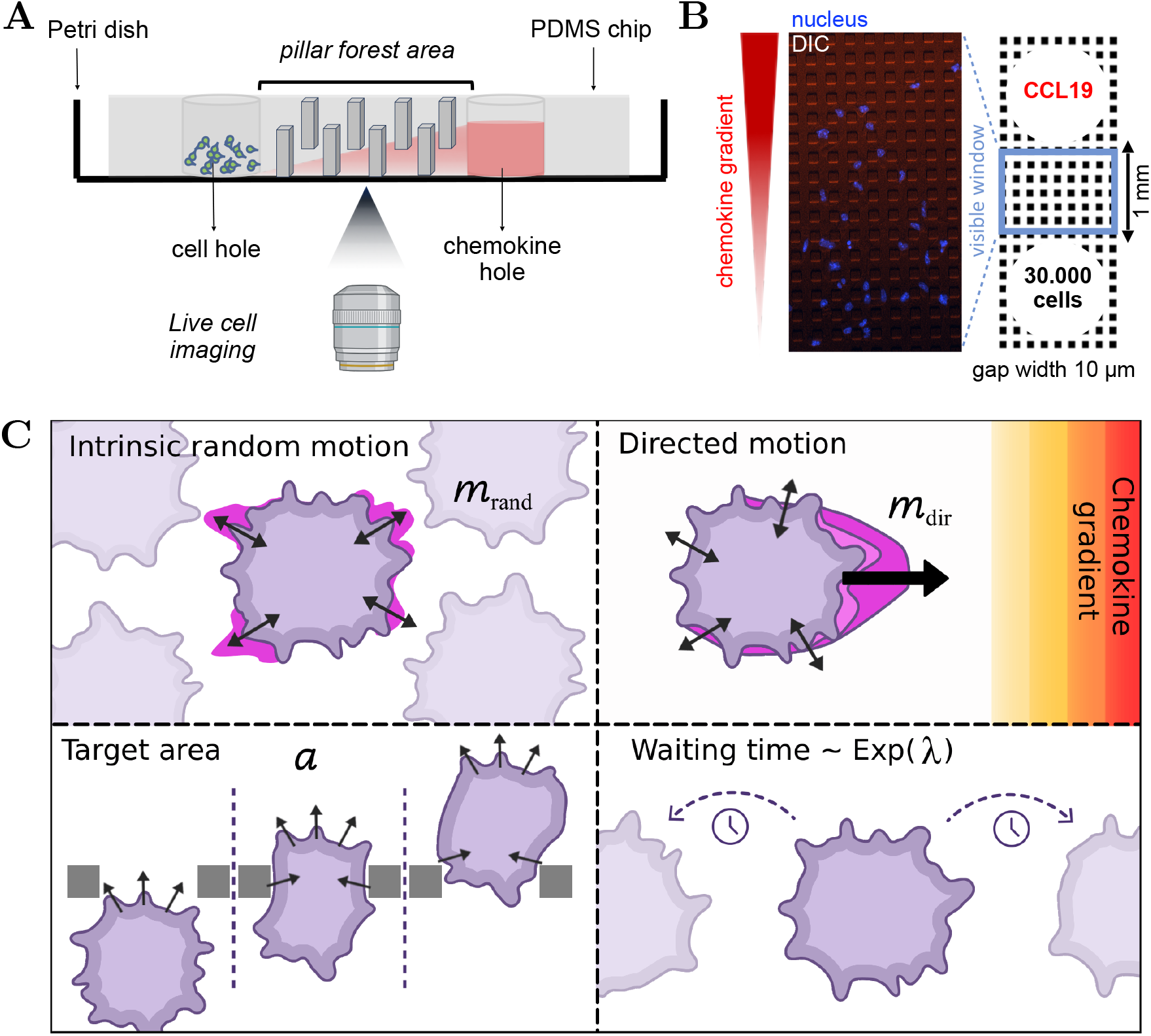
Schematic depiction of the experimental and modeling setup. (**A**) Dendritic cells are placed in an entry hole on the left and follow a chemokine gradient to the right through the pillar forest on a microstructured polydimethylsiloxane (PDMS) chip. (**B**) Top-down differential interference contrast (DIC) microscopy view within the visible window of the pillar forest, where cell movement can be tracked. Dextran was added to the CCL19 solution to visualize the chemokine gradient. (**C**) Visualization of a cell in the Cellular Potts model and the four model parameters to be estimated: the intrinsic random motion strength *m*_rand_ (upper left tile), the chemokine attraction strength *m*_dir_ (upper right tile), the target area of a cell *a* (lower left tile), and the rate *λ* until the cell changes its intrinsic direction (lower right tile).

From these trajectories, we extracted descriptive measures of migration that captured complementary aspects of motility (see Methods). Displacement quantifies overall progress across the track, velocity reflects the average speed of migration, and turning angle and angle degree characterize directional biases toward the chemokine gradient or deflections caused by obstacles.

These summary statistics form the basis for quantitative comparison with a mechanistic model, linking experimental single-cell measurements to simulations and providing insight into how dendritic cells balance chemokine guidance with geometric restrictions imposed by tissue-like barriers.

### Modeling cell migration dynamics in complex spatial environments

To analyze dendritic cell migration, we developed a 2D Cellular Potts model with chemotactic signaling and persistent random motion embedded in a simulation-based inference pipeline. In this lattice-based stochastic framework, cells are represented as extended domains of lattice sites, allowing the simulation of realistic cell shapes and their deformations. Cell behavior emerges from an energy minimization principle that combines terms for volume constraints with responses to external cues, such as chemokine gradients, allowing us to generate trajectories under conditions that mimic the experimental setup (Figure 1C).

The model possesses four parameters with a substantial influence on the migration patterns: the chemokine attraction strength *m*_dir_, which quantifies the sensitivity of cells to the chemokine gradient modulating directed movement; the intrinsic random motion strength *m*_rand_, which governs the bias toward previously randomly chosen movement directions; the rate *λ* for the waiting time distribution between directional changes of the intrinsic random motion; and the target area *a*, which regulates cell size via area constraints (for details, see Methods).

To estimate these parameters from data, we established a simulation-based inference pipeline. This pipeline supports (i) approximate Bayesian computation (ABC) with hand-crafted summary statistics, (ii) ABC with neural feature extractors trained to predict posterior means (ABC-PM), (iii) ABC with inference-tailored summaries (ABC-NPE), and (iv) neural posterior estimation (NPE) using jointly trained summary networks (see Methods). For NPE, we trained a summary network jointly with an inference network on a training dataset of 32,000 simulation–parameter pairs using an additional 100 simulations for validation. The inference network was implemented as a conditional normalizing flow optimized by minimizing the Kullback–Leibler divergence. The learned summary network is also used in ABCNPE and can alternatively be adapted to directly predict the posterior mean by optimizing a mean-squared error objective without the inference network.

This pipeline enables inference of Cellular Potts model parameters from dendritic cell trajectories, linking experimental migration data to the underlying stochastic processes shaped by chemokine gradients and physical barriers.

### Inference-tailored summaries enable accurate inference on simulated cell movement in complex environments

To infer the parameters of the cell migration model, we established a parameter estimation pipeline. To ensure reproducibility, we conducted experiments using three synthetic datasets generated by the simulation model. For each dataset, we attempted to recover the underlying ground truth parameters, which were sampled from a prior distribution.

The ABC methods were run until either a minimum acceptance rate of 0.01 was reached or a maximum of 15 sequential generations was completed, including a pre-calibration round. In total, this required up to 432,000 simulations and runtimes between 2 and 20 h (Table 1). The assessment of the inference results revealed that all ABC approaches converged, as indicated by the low acceptance thresholds in the last generation (Supplementary Figure 1–3). For NPE, the inference procedure is amortized, meaning that once the networks are trained, the parameters can be inferred for any new dataset without retraining. To assess the convergence and calibration of the posterior in this setting, we applied simulation-based calibration on a larger validation dataset of 100 simulations, which confirmed good calibration and showed no visible bias (Supplementary Figure 4). Such rigorous validation was not feasible for the ABC methods due to their prohibitive inference time.

**Table 1:** Comparison of the different inference methods on simulation budget and performance metrics. ABC-based methods were evaluated on three representative simulated datasets, whereas NPE was validated on 100 datasets. We computed the normalized root mean squared error (NRMSE). The training time accounts for both training set generation and neural network training and was measured on a high-performance computing cluster (for details, see Methods).

| Method | # Simulations | Training | Inference | # Generations | NRMSE |
| --- | --- | --- | --- | --- | --- |
| ABC | 66,000–413,000 | – | 2.1–19.2 h | 12–16 | 0.23 |
| ABC-PM | 59,000–432,000 | 1.5 h | 1.8–12.2 h | 11–16 | 0.48 |
| ABC-NPE | 120,000–428,000 | 4.0 h | 3.7–13.4 h | 13–16 | 0.23 |
| NPE | 32,100 | 4.0 h | 1 s | – | 0.20 |

In terms of accuracy, measured via the normalized root mean squared error (NRMSE), NPE yielded the smallest error, closely followed by ABC and ABCNPE (Table 1). Interestingly, the neural network trained on the posterior mean achieved a smaller NRMSE of 0.17 when directly predicting the parameters on the validation data. However, when used as a feature extractor in the ABC-PM, the error was much higher, with an NRMSE of 0.48 on the three synthetic datasets. In contrast, NPE achieved better performance using far less simulations with a training duration of approximately 4 h (Table 1). Moreover, inference with NPE is essentially instantaneous because the networks are trained once and can then be readily applied to new data, enabling the efficient processing of large numbers of datasets and allowing us to analyze parameter identifiability. For instance, the identifiability assessment for the rate parameter *λ* of the waiting time revealed an overall lower posterior contraction (Supplementary Figure 5) and a diminished parameter recovery accuracy with NPE on the validation data (Supplementary Figure 4). This lower identifiability may be attributed to the temporal resolution of the observations (30 s between each image), which limits the amount of informative signal available for estimating this parameter.

Our results suggest that the ABC method, which relies on the posterior mean, exhibits a systematic bias in one of the parameters: the true value of the intrinsic random motion strength *m*_rand_ frequently lies outside the high-density region of the inferred posterior distribution (Supplementary Figure 5), despite the summary network itself showing no apparent bias during validation (Supplementary Figure 4). In contrast, the use of hand-crafted or inference-tailored summaries led to improved parameter recovery in ABC (Table 1).

An exploratory analysis of latent summary representations reveals that one advantage of posterior mean summaries lies in their interpretability: each component corresponds directly to a model parameter, rendering the representation transparent and easily analyzable, whereas the latent summary representations learned jointly with the normalizing flow are not inherently interpretable. However, the learned summaries in NPE were correlated with specific model parameters, indicating a degree of functional disentanglement in the learned representation (Supplementary Figure 6). Moreover, a simple regression of the latent dimension onto the parameters already allows the prediction of the true parameters (Figure 2A).

**Figure 2:**
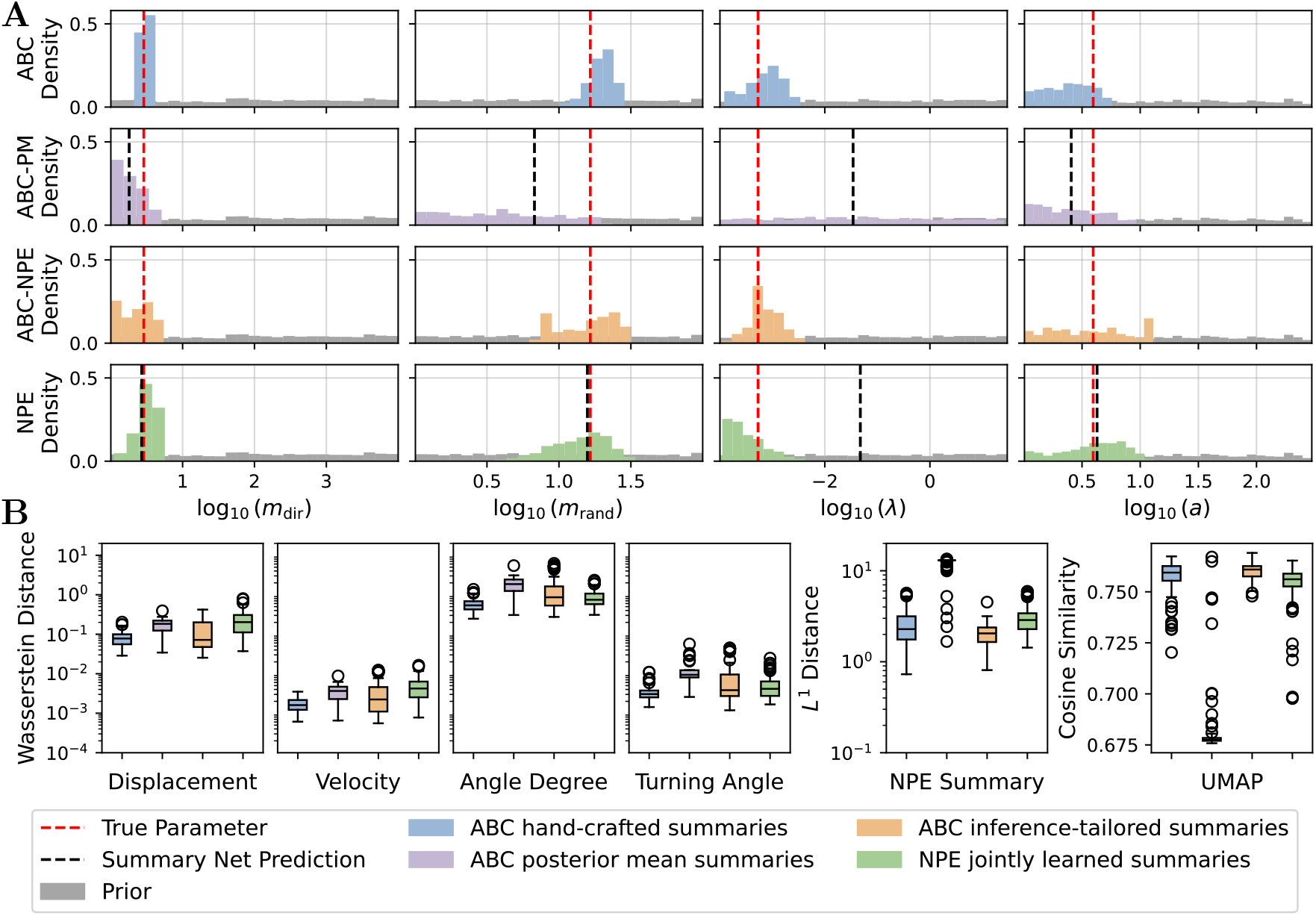
Posterior approximations and summary statistics on synthetic data. (**A**) Posterior approximations using the different ABC approaches and NPE with jointly trained summaries and predictions from summary latent spaces. (**B**) The distance between hand-crafted summaries and the statistics of the synthetic test set was quantified using the Wasserstein distance, the *L*^1^ distance computed in NPE summary space, and cosine similarity in UMAP representation. All metrics were based on 100 simulation samples drawn from each posterior. Distributions are shown as box plots, indicating the medians, quartiles, and outliers.

To better understand these findings, we examined the informativeness of the summary statistics across methods. Specifically, we computed the hand-crafted summaries for simulations drawn from all four inferred posteriors. If a cell was not observed in the simulation because it moved outside the visible window, it was not included in the corresponding statistics. When using the metric employed by ABC, the Wasserstein distance, simulations from the posteriors and synthetic test data were similar across all approaches, despite their distinct posterior distributions (Figure 2B). However, alternative distance metrics, such as the *L*^1^ distance in the NPE summary space and cosine similarity on a UMAP projection, reveal notable deviations, particularly between ABC with posterior mean summaries and the other methods. This suggests that the hand-crafted summaries may be insufficiently expressive to capture the full range of informative variations of the stochastic model.

In summary, although hand-crafted and posterior mean summaries remain interpretable, they can fail to capture the variability required for reliable parameter recovery. In contrast, inference-tailored summaries adapt their representation to the data and inference task, leading to improved parameter recovery. By optimizing the summary space directly for parameter estimation, this provides a flexible and data-driven approach that enables more accurate and robust inference.

### Robust model calibration and validation of cellular dynamics on empirical data

Given the validated inference pipeline, we investigated the interplay between chemokine signaling and physical constraints in dendritic cell migration using experimental data (Figure 3A). The dataset contained 143 individual cell trajectories of varying lengths (18 to 120 frames per cell, with a mean duration of 33.2 min), reflecting heterogeneity in tracking durations across the experiment.

**Figure 3:**
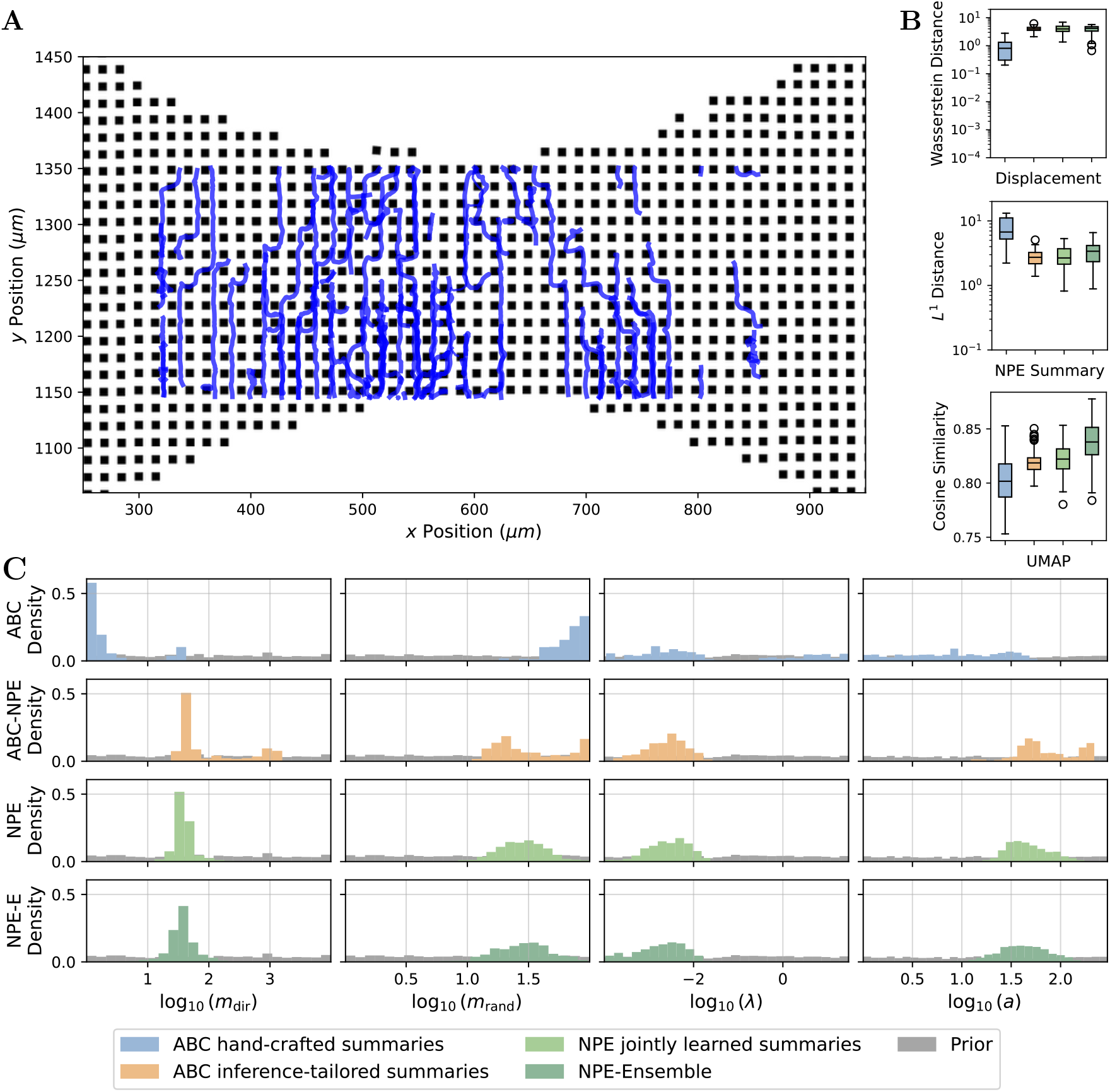
Posterior approximations and summary statistics on experimental data. **(A)** Experimental data (cells in blue) within the pillar forest are shown. (**B**) Distances between the hand-crafted summaries in the NPE summary space and UMAP representation were computed. All metrics are based on 100 simulation samples drawn from each posterior. Distributions are shown as box plots, indicating medians, quartiles, and outliers. (**C**) Posterior approximations using different ABC approaches and NPE with jointly trained summaries.

ABC was performed using both hand-crafted and inference-tailored summary statistics. Both approaches converged after 13 generations, as indicated by the small final acceptance thresholds, requiring approximately 200,000–300,000 simulations and 27–50 h of runtime (Supplementary Figure 7). In addition, NPE was employed, including an ensemble of three independently trained networks (NPE-E), to obtain a more robust estimate that is less sensitive to variations in individual network training.

To assess the differences between the approaches, we compared the hand-crafted summary statistics on posterior simulations, the *L*^1^ distance in the NPE summary space, and the cosine similarity in the UMAP projection. Among the hand-crafted summaries, only the displacement feature shows a noticeably smaller distance for ABC with hand-crafted summaries (Figure 3B); the remaining summaries yield highly similar distances across all methods (Supplementary Figure 8). In contrast, the *L*^1^ distance and cosine similarity metrics indicate that the predictions from the NPE-based methods are generally closer to the observed data. The ensemble NPE achieved the highest cosine similarity, whereas both the single and ensemble NPEs exhibited nearly identical *L*^1^ distances. Because the UMAP projection was not used during training and inference in any method, this metric offers an independent measure of posterior predictive quality. As an additional robustness check, we repeated the inference with NPE on parts of the dataset and controlled for model misspecification (see Supplementary Material).

The posterior distributions obtained from the ensemble and single NPE models were highly similar, whereas the ensemble was more conservative, indicating consistency across the training runs (Figure 3C). In contrast, the inference-tailored ABC posterior samples show slightly greater uncertainty and hint at a potentially multi-modal posterior, although its median aligns with that of the NPE-based estimates (Figure 3C). Notably, ABC using hand-crafted summaries yielded a markedly different posterior, with a distinct shift in the median and signs of bimodality (Figure 3C). For instance, the smaller mode in the inferred chemokine attraction strength *m*_dir_ aligns with the dominant mode found in the NPE posteriors (Figure 3C).

The median estimated cell area from NPE-E is *a* = 41.73 µm^2^ (Figure 3C), aligning with the expected diameter of 10–15 µm [44, BNID 113239]. Although the model does not make any specific assumptions regarding the cell shape, this area suggests that the cells are larger than the gap between the two pillars (10 µm). Additionally, the median estimated rate parameter is *λ* = 2.1 ms^−1^, corresponding to a mean waiting time of 7.9 min, which is more than one order of magnitude larger than the time of 30 s between two images.

In conclusion, our results demonstrate the robustness and flexibility of our Cellular Potts model in capturing key cellular dynamics. By integrating posterior inference with neural summaries, we successfully identified critical parameters that govern cell migration.

### Simulation study shows modulating role of physical obstacles and persistence in cell migration

To assess how physical structures modulate migratory behavior, we used our computational model and inference pipeline to investigate the impact of obstacles on cell movement. Specifically, we aimed to disentangle the independent contributions of chemokine guidance and persistence in random motility.

Based on the posterior median of the model parameters from the NPE ensemble, we simulated cell migration under four distinct conditions: with (Figure 4A) and without a chemokine gradient that was constant over time (Figure 4B), with an initial chemokine signal lasting only 30 min (Figure 4C), and with suppressed persistence in random movement (intrinsic random movement was set to *m*_rand_ = 0) but a constant chemokine gradient (Figure 4D). Simulations were sampled at a higher resolution of 10-second intervals and also recorded outside the visible window of the experimental setting, enabling controlled comparisons of cell responses to environmental cues influencing migration dynamics.

**Figure 4:**
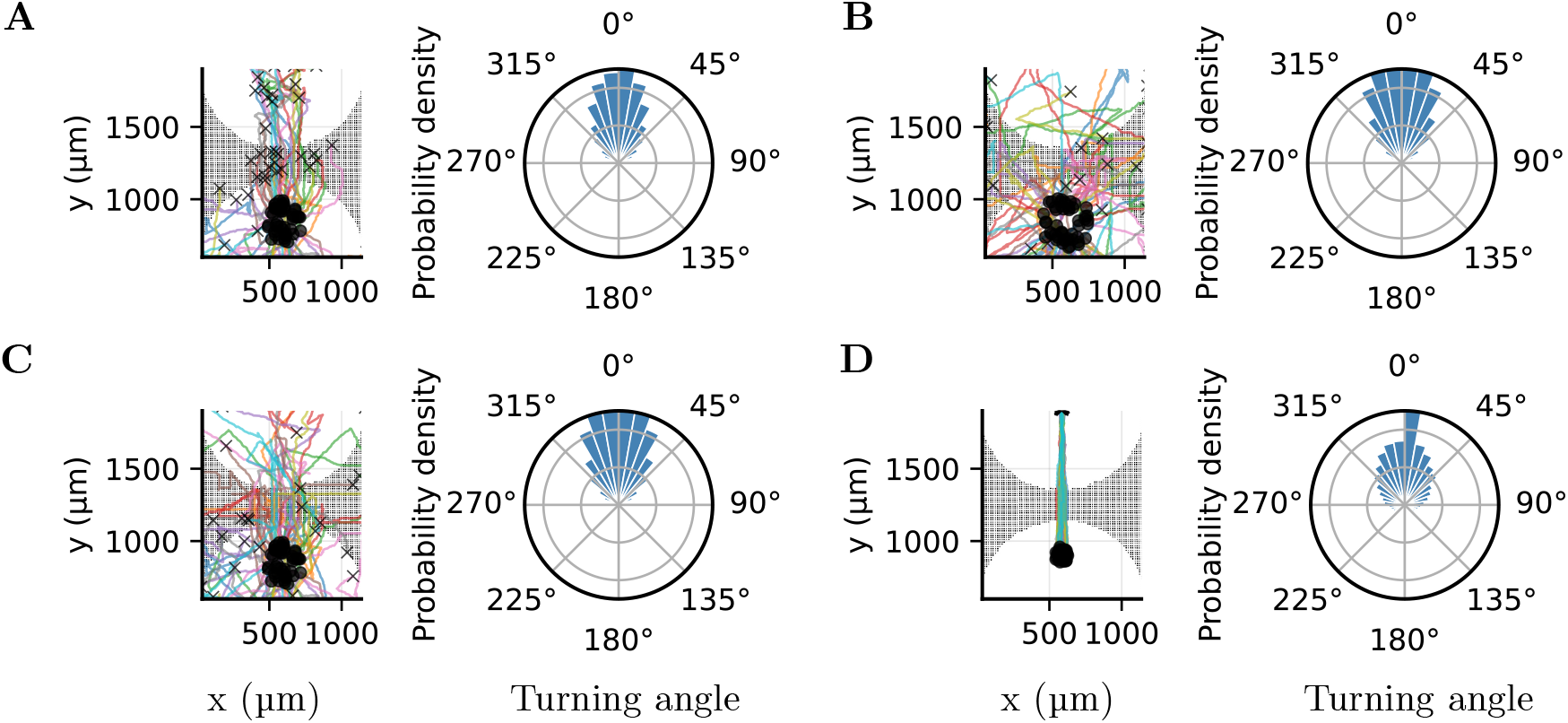
Simulation study shows modulating role of obstacles. The trajectory of 50 random cells (left) and the distribution of the turning angles (of all cells) (right) are shown. The endpoints are marked with ‘x’ if visible in the selected area. (**A**) Simulation setup mirroring the experimental setup with structural obstacles and a constant chemokine gradient. (**B**) Simulation setup without a chemokine. (**C**) Simulation setup in which the chemokine is active only for 30 min. (**D**) Simulation setup without persistence in the random walk but with a constant chemokine.

The results demonstrate that persistence is a key driver of cell dispersion (Figure 4). Even in the absence of directional guidance through the chemokine gradient, cells with persistent movement spread widely throughout the environment (Figure 4B). In contrast, when persistence is removed, migration is spatially constrained and directionally limited (Figure 4D). The pillar forest further modulates this behavior by imposing spatial constraints that can trap cells. Moreover, simulations with a short-lived chemokine gradient revealed that the removal of guidance cues allowed trapped cells to reorient and resume dispersal (Figure 4C). This suggests a dynamic interplay: while chemokines bias movement, their prolonged presence may restrict escape from local traps in obstacle-rich environments.

This phenomenon became even more apparent when analyzing the number of cells that reached the upper clearing of our experimental setup. Simulations where the chemokine signal is active only at the first 50–75 % of the simulation time show a higher proportion of successful arrivals compared to scenarios with constant guidance (Supplementary Figure 9). Our experimental setup does not allow us to directly observe and thus verify this outcome, as our imaging is limited to the pillar forest region and does not extend to the final source location. Nevertheless, a similar effect may occur *in vivo*, potentially because of signal decay or attenuation near the source. This counterintuitive result highlights that temporally modulated guidance cues can facilitate more efficient exploration in the presence of obstacles.

Further analysis revealed that persistence is particularly important for enabling movement orthogonal to the chemokine gradient (Figure 4D). These findings underscore that intrinsic motility parameters and environmental features jointly determine the range and structure of cellular trajectories.

In summary, the simulations show that physical obstacles actively reshape migration, not only by slowing cells but also by altering the directionality and dispersion of their paths. While chemokines impose a directional bias, obstacles introduce nonlinear constraints that modify or obscure this signal in the observable data.

## Discussion

A quantitative understanding of cell migration is essential across diverse domains, including immunology, oncology, and tissue engineering, where directed cell movement plays a central role in physiological and pathological processes. Despite its importance, methods for inferring the parameters of mechanistic models of migration from experimental data remain limited, especially in spatially structured or noisy microenvironments, where hand-crafted summaries may fail to capture the relevant dynamics.

The findings of this study provide a foundation for researchers to enhance modeling approaches for biological systems. Specifically, our study underscores the substantial impact of spatial obstacles on cell migration, as evidenced by the manner in which the pillar forest environment regulates both the speed and directional alterations. Additionally, we demonstrate that the learned summaries exhibit superior performance compared to traditional summary statistics in the context of simulation-based inference. Our simulation study further revealed that persistent cell motion plays a critical role in overcoming spatial constraints and that temporally modulated chemokine cues can facilitate dispersal in obstacle-rich environments.

The pillar forest used in our simulations provides a well-controlled and interpretable setting for uncovering the fundamental principles of cell migration. Building on our framework, future studies could examine how cells behave in spatially and temporally complex environments, where guidance cues may be transient, competing, or context-dependent. It remains an open question how cells resolve directional conflicts in the presence of multiple chemotactic sources or how robust their migratory behavior is in the face of spatial noise or fragmented gradients. Addressing these challenges would advance our understanding of how immune cells navigate physiologically realistic environments and flexibly respond to complex environmental stimuli.

The utilization of learned summaries in ABC shares similarities with other approaches that employ an estimator trained on pairs of simulations and parameters to predict the posterior mean or learn a summary based on an autoencoder [40, 45]. Yet, the latent space of these autoencoders is not specifically tailored for inference but for reconstruction of the data. The posterior mean estimators are then used in the context of ABC to generate summary statistics [37, 46, 47, 48, 49]. However, we demonstrated that the posterior mean was not sufficiently informative. Our results suggest that ABC-PM exhibits a systematic bias, despite the summary network itself showing no apparent bias during validation. This discrepancy likely arises because predicting only the posterior mean does not adequately capture the uncertainty inherent in the data, resulting in biased estimates. Therefore, it was recently proposed to append noise estimates or higher-order moments of the parameters to the posterior mean [40, 39]. However, this approach remains a type of manual feature engineering. Moreover, although the neural network architecture is identical for both the posterior mean and inference-tailored summaries, joint training with the normalizing flow removes the constraint that the output dimensionality of the summary network matches the parameter space, thereby enabling greater representational flexibility.

Adaptive distances based on hand-crafted summaries have been extensively studied [38, 39]. Yet, identifying the optimal distance measure depends heavily on the informativeness of the chosen statistics, which may fail to capture the necessary information. In contrast, the joint learning of summaries with a neural posterior estimator ensures that the summaries are maximally informative and eliminates the need for additional upfront simulations. Moreover, our findings indicate that ABC requires 10 times more simulations to attain comparable accuracy, thereby highlighting the efficacy of neural posterior estimation, which is consistent with previous findings [50, 51]. As cells are heterogeneous in size and even have specific shapes depending on the chemical gradient [52], a further direction could be to quantify individual differences between cells using an amortized hierarchical framework [53, 54].

Instead of applying the Wasserstein distance to the hand-crafted summaries, the distance has been utilized directly on the observed data [55]. However, this approach has not been generalized to a spatial context with a varying number of observations and therefore cannot be applied in our context. In general, if the quantiles of the statistic (or the data) do not vary significantly as the parameter changes, the Wasserstein distance will not change in a meaningful manner and thus cannot serve as an accurate posterior approximation [56].

An opportunity to extend our model lies in its current assumption of a homogeneous and static environment. While this simplification has provided clear insights into core migration behaviors, it does not capture the dynamic, feedback-rich nature of many biological systems, where cells actively remodel their surroundings. For example, the chemokine concentration diffuses over time, migrating cells may reshape local chemotactic gradients [57, 58], restructure the extracellular matrix [59], or exert mechanical forces, establishing feedback loops that influence both their own motility and collective behavior [60]. Moreover, dendritic cell migration is tightly integrated with their immunological functions, such as antigen capture, cytokine production, and T-cell priming, and is influenced by a complex interplay of extracellular cues and intracellular signaling pathways [61]. Incorporating such bidirectional cell–environment interactions into future models offers the potential for more faithful representations of the adaptive and co-evolving dynamics that characterize living tissues. Experimental evidence further shows that dendritic cells require stable spatial chemokine gradients to maintain directional persistence, as transient or oscillatory cues rapidly disrupt cytoskeletal polarization and migration [62]. This highlights that dynamic environmental signals may not only fail to support chemotaxis but can actively destabilize it, reinforcing the need to jointly model spatial and temporal features.

The findings reveal that learned summaries, tailored specifically for inference, exhibit superior performance compared to hand-crafted summary statistics and provide more information than summaries based on the posterior mean. This enhancement in performance can be attributed to the adaptability of neural networks, which automatically extract relevant features from data, thereby eliminating the need for manually designed summaries. Instead, these networks learn features that are specifically relevant to the inference task. By leveraging this approach, researchers can develop more complex models and calibrate them effectively with experimental data, paving the way for deeper insights into the fundamental mechanisms of cell migration.

## Methods

In this study, we analyzed the movement of bone marrow-derived dendritic cells (DCs) in complex microenvironments, where both chemical signaling and physical barriers influence their movement.

### Experimental procedures

DCs are confined in an environment with 7 ng of the chemokine (C-C motif) ligand 19 (CCL19), which binds to the C-C chemokine receptor type 7 (CCR7) expressed on the surface of mature DCs [63]. To create physical obstacles, the area between the two confining surfaces was intersected by pillars with defined gaps: a “pillar forest” [43] (see Figure 1). The presence of these obstacles introduces physical complexity, compelling cells to traverse the gaps of 10 µm. This adds a layer of mechanical challenge to their movement, as the cells constantly encounter pillars on their migratory path.

### Dendritic cell culture from mouse bone marrow

Femurs and tibias from the legs of 6–12 week-old C57BL/6J mice were removed and placed in 70 % ethanol for 2 min. Bone marrow was flushed with PBS using a 26-gauge needle. 2 × 10^6^ cells were seeded per 100 mm Petri dish containing 9 mL of complete medium (Roswell Park Memorial Institute (RPMI) 1640 supplemented with 10 % Fetal Calf Serum (FCS), 2 mM L-Glutamine, 100 U/mL Penicillin, 100 µg mL^−1^ Streptomycin, and 50 µM *β*-mercaptoethanol; ThermoFischer Scientific) and 1 mL Granulocyte–Monocyte Colony Stimulating Factor (GM-CSF, supernatant from hybridoma culture). On days 3 and 6, complete medium supplemented with 20 % GM-CSF was added to each dish. To induce maturation, the cells were stimulated overnight with 200 ng mL^−1^ lipopolysaccharide (LPS) and used for experiments on day 8–9. Alternatively, DCs were frozen in FCS containing 10 % DMSO on day 7. For experimental use, the cells were thawed the day before the experiment and stimulated with 200 ng mL^−1^ LPS overnight.

### Photomask patterning

Photomasks were drawn in CorelDraw and converted to Gerber in LinkCad. Photomasks were fabricated at JD Photodata (UK). These photomasks were used to define microscale patterns on a photoresist-coated silicon wafer during UV lithography. The transparent and opaque regions of the mask control the areas where UV light reaches the photoresist, enabling precise pattern transfer. To produce the master wafer, a 4 inch wafer was first baked at 110 °C for 5 min. 6005 TF SU8 was spun up to 3000 RPM with an acceleration of 300 RPM/s for 30 s. The wafer was then baked at 110 °C for 5 min. The wafer was exposed to 100 mJ/cm^2^ UV light through the photomask, then baked for 5 min at 110 °C, and then developed in SU8 developer. As a final step, the wafer was baked at 135 °C for 5 min to permanently harden the SU8. The height of the wafer was approximately 5 µm.

### Pillar forest migration assay

Microfabricated chips were produced by pouring a polydimethylsiloxane (PDMS) mixture (silicon rubber and curing agent at a ratio of 10:1 by weight) onto a silicon wafer with a previously designed photomask pattern. This was incubated at 80 °C overnight after removing air bubbles by placing the PDMS in a vacuum desiccator. The PDMS was separated from the wafer, and single chips were cut out using a surgical blade. Next, two holes (diameter of 1.5 mm) on each side of the pillar forest area were punched at a distance of 1 mm, and the PDMS was bound upside-down to a glass-bottom dish. To activate the surface, both the PDMS chip and dish were cleaned with plasma before assembly. To equilibrate the chip, it was flushed with complete medium and incubated at 37 °C, 5 % CO_2_ for at least 1 h. Afterward the medium was removed from the two holes, and they were filled with 7 µL CCL19 solution at a concentration of 1 g mL^−1^ on one side and 3 × 10^4^ cells in complete medium on the opposite side. Additionally, dextran was added to the CCL19 solution (1:1000) to visualize the chemokine gradient. BMDCs were stained with NucBlue (Invitrogen) and concentrated to 6.25 × 10^6^ cells/mL in advance.

### Microscopy and image analysis

After a 2–2.5 h incubation at 37 °C, 5 % CO_2_, samples were imaged with an inverted wide-field Nikon Eclipse Ti-2E microscope in a humidified and heated chamber at 37 °C and 5 % CO_2_ (Ibidi Gas Mixer), equipped with a Plan-Apochromat 20 ×/0.8 air objective, a DS-Qi2 camera, and a Lumencor Spectra X light source (390 nm, 475 nm, 542/575 nm; Lumencor). Movies were taken for 2 h, recording one image every 30 s. Videos were analyzed using `Fiji` [64]. Cell nuclei tracking was performed manually or automatically using the `Fiji` plugin `TrackMate` [65]. However, owing to the microscope setup, tracking was only possible within the pillar forest between the two clearings.

### Model description

Cell migration behavior was described using a modified Cellular Potts model (CPM), a lattice-based framework that effectively represents both individual cell dynamics and interactions within complex environments [17]. Each cell is represented as a set of connected lattice sites on a discrete 3-dimensional grid. The system evolves through stochastic updates that minimize the overall energy function *H* using a Metropolis algorithm. In each iteration, a lattice site *i* and one of its neighboring sites *j* are randomly selected. Cell *j* is proposed to replace cell *i*, and the resulting change in energy Δ*H* is calculated. The proposed change is then probabilistically accepted based on the Metropolis criterion.

The CPM energy function is given by

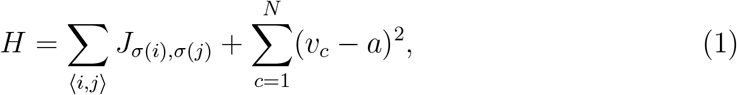

where *σ*(*i*) denotes the cell at site *i*, and the first sum runs over neighboring lattice sites ⟨*i, j*⟩ with *σ*(*i*) ≠ *σ*(*j*). The interaction term *J*_*σ*(*i*),*σ*(*j*)_ models the interface energies between different cells and the medium. The second term penalizes deviations of a cell’s volume *v*_*c*_ from a target volume *a* and contributes to realistic cell shapes. This volume constraint indirectly affects cell motility by limiting the number of lattice sites a cell may occupy. The combined effects of surface tension and volume conservation allow the model to capture mechanical interactions with neighboring cells and the extracellular matrix. In dense environments, deviations from the preferred volume can generate mechanical pressure, resulting in tissue-level reorganization.

To model chemotactically guided cell migration, we extended the basic CPM with two additional components: directional movement in response to chemokine gradients and intrinsic persistent random motion. To bias cell movement toward higher concentrations, the chemotactic signal is integrated into the energy function via an additive term

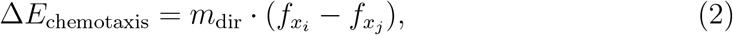

where 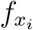 and 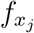 are the local chemoattractant concentrations at the occupied and empty lattice sites, respectively, and *m*_dir_ controls the strength of chemotactic attraction [66]. The update is energetically favored if the move leads to a position with a higher concentration.

To account for undirected and exploratory cell movements, we implemented a persistent random walk. This implementation resembles a run-and-tumble process and reflects experimentally observed persistent random motion [22]. Each cell was assigned a preferred migration direction that was periodically updated. The energy contribution for a proposed move in the direction ***s***_*c*_ is given by another additive term to the energy function *H*:

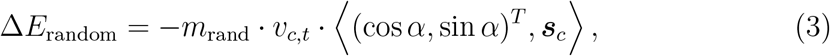

where *α* is the current migration angle, *v*_*c,t*_ is the current area of the cell *c*, and *m*_rand_ is the strength of intrinsic motility. The scalar product penalizes deviations from the preferred direction. For each cell, the angle *α* is updated after a waiting time *t*_*α*_ ∼ Exp(*λ*) with rate *λ*, and a new direction is sampled uniformly from 𝒰 [0, 2*π*].

Because the cells are comparatively flat in the considered experimental setup, a 2D model was used to capture the exploration details without adding unnecessary complexity. We used an image to position a dense array of pillars between the two ends of the migration area, thereby reproducing the spatial constraints of our experimental setup. The model was built using `MorpheusML`, an XML format for encoding multiscale models and multicellular biological systems [22]. Spatial resolution is set to 1.31 µm per node length. To emulate the experimental setting, we simulated cell movement in the entire area and extracted the coordinates of cells in the pillar forest at 30-second intervals. The chemokine gradient was modeled as a time-independent Gaussian distribution with a peak concentration of 7 × 10^6^ µm^−2^ and a standard deviation of 550 µm. This choice reflects typical diffusion distances and ensures a smooth and biologically meaningful gradient.

In summary, the parameters to be inferred from the data in this study are

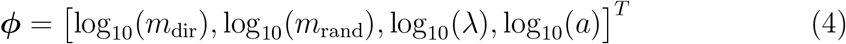

with their corresponding priors provided in Table 2. Uniform priors were specified on the logarithmic scale, and bounds were selected based on biological constraints or the computational feasibility of the simulations. The feasibility of the simulations was assessed by evaluating the simulations at the upper and lower bounds of the parameter space. Additional technical constants used in the CPM are documented in the model file, provided here: https://identifiers.org/morpheus/M6342.

**Table 2:** Parameters to estimate in the cell migration model. Parameters were transformed to a logarithmic scale and had uniform priors.

| Parameter | Description | Prior |
| --- | --- | --- |
| $\log_{10}(m_{\text{dir}})$ | Influence of chemokine attraction | $\mathcal{U}[0, 4]$ |
| $\log_{10}(m_{\text{rand}})$ | Strength of intrinsic random movement | $\mathcal{U}[0, 2]$ |
| $\log_{10}(\lambda)$ | Rate of the waiting time distribution ($\text{s}^{-1}$) | $\mathcal{U}[-4, 1.5]$ |
| $\log_{10}(a)$ | Area of cells ($\mu\text{m}^2$) | $\mathcal{U}[0, 2.5]$ |

### Hand-crafted summary statistics

The dynamics of the model depend on several unknown parameters that must be estimated from the available data. We employ approximate Bayesian computation using hand-crafted summary statistics that were built specifically for this model, referred to as *ABC with hand-crafted summary statistics*.

Each simulation contains multiple cell tracks of varying lengths, as cells could move out of the observation area or leave the 2D plane. Because the trajectory of a cell is governed by stochastic movement, direct matching of a simulated cell to a real cell is not possible. Instead, summary statistics that characterize individual cell movement were constructed. These statistics were computed for each cell, and the distributions of these statistics were compared between the simulation and observed data. This approach eliminates the need for direct track matching and allows for flexibility in the number of simulated cells. However, for reliable parameter inference, the number of cells should not deviate too much, as crowded and sparse environments differ significantly; for example, cells may block each other more frequently in a crowded environment.

We constructed four different widely used summary statistics to describe different aspects of cell movement [67, 68]. Denoting the position of a cell *c* ∈ {1, …, *N*} as 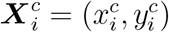 at time step *i* for *i* ∈ {1, …, *T* ^*c*^}, we define the following statistical quantities. Note that the time steps of each cell are relative to the first time they were observed.

- **Displacement (D)** measures the squared distance traveled by the cell *c* over the observed time:

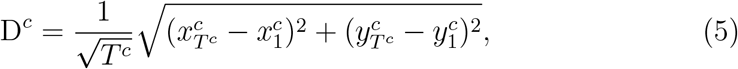

where *T* ^*c*^ is the number of observed time steps for each cell. We scaled the displacement by time so that different observed sequence lengths could be handled. For a standard random walk, the expected translation distance after *T* steps is of the order of 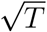.
- **Velocity (V)** is the average speed of a cell *c* and is expressed as the mean over all time steps

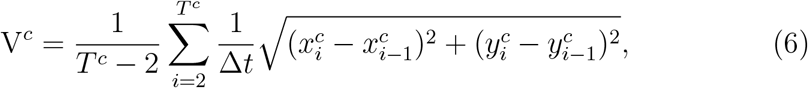

where Δ*t* = 30 s is the interval at which the cells were observed.
- **Turning angle (TA)** quantifies the mean change in directionality of a cell *c* between consecutive movements:

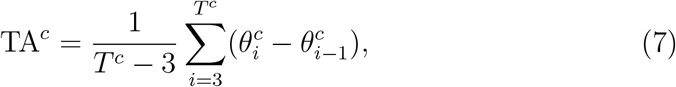

where 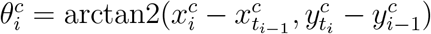.
- **Angle degree (AD)** captures the average absolute direction between two consecutive positions:

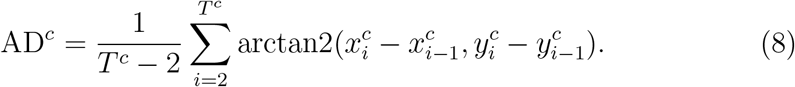

To compute the acceptance criterion in ABC, we used the Wasserstein distance to calculate the distance between the samples of the distributions of the summaries of the simulations and the summaries of the experimental data. The Wasserstein distance quantifies the cost of transforming one distribution into another, thereby providing a comprehensive measure of discrepancy (see [69] for a detailed introduction). By comparing the distributions of the summary statistics for the simulated and observed cells, we captured the differences across the entire dataset, avoiding the matching of trajectories. To integrate multiple statistics while minimizing the overreliance on any single summary, the Wasserstein distance was computed separately for each statistic. To normalize the contribution of each summary, the distance is weighted by the inverse of the difference between the largest and smallest statistics, as suggested in [70]. These weights were calculated from simulations of a pre-calibration population with parameters sampled from the prior. Finally, the weighted distances are aggregated by summing them up.

### Neural posterior estimation with learned summary statistics

Neural posterior estimation (NPE) methods leverage neural networks to learn mappings from data to model parameters and can integrate learned summary networks that automatically extract relevant features from the data [42]. Instead of relying on hand-crafted summary statistics, we used a summary network *s*_***ψ***_ with trainable parameters ***ψ***, designed to handle variable-length inputs, both in terms of the number of observed time points per cell and the number of cells per experiment. To extract informative features from each individual cell trajectory, we applied a combination of one-dimensional convolutional layers and a recurrent neural network, which is a standard architecture for time-series processing. To aggregate across multiple cells, we employed an attention-based pooling mechanism that produced a fixed-length summary vector while preserving flexibility with respect to the cell count (see Implementation for details). The summary network is then utilized to learn a *d*-dimensional representation *s*_***ψ***_(***X***) ∈ ℝ^*d*^ of the observed data points 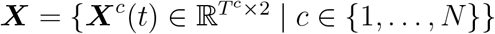.

The summary statistics serve as inputs to a conditional normalizing flow, which transforms the posterior into a simpler density from which we know how to sample. This method allows efficient and accurate sampling and density evaluation [71, 72]. Let ***z*** be a latent variable described by a multivariate normal distribution *p*(***z***). The model parameters ***ϕ*** are mapped to these latent variables conditional on the data *s*_***ψ***_(***X***) by a normalizing flow 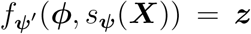. This invertible transformation is parameterized by ***ψ***^′^ and, by construction, has a tractable Jacobian. The approximation 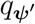 to the target density *p*(***ϕ*** | *s*_***ψ***_(***X***)) is given by the change-of-variables formula

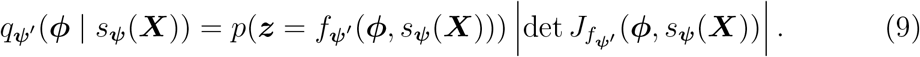

If we know 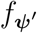, we can sample from the posterior by sampling ***z*** ∼ *p*(***z***) and applying 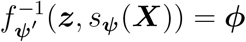.

To train the summary network *s*_***ψ***_ jointly with the conditional normalizing flow 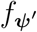, we follow the work of [73] and minimize the Kullback-Leibler divergence between the true and approximate posterior distributions:

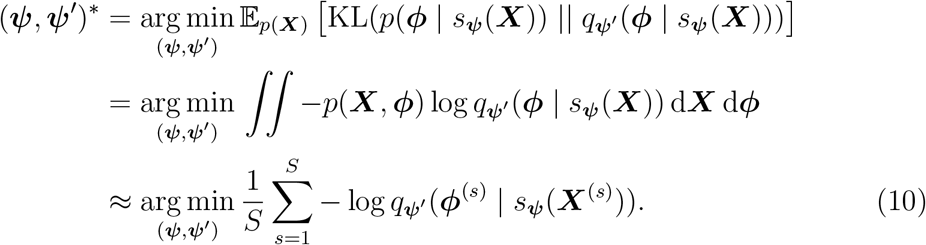

The integral is approximated using *S* samples from the joint distribution *p*(***X, ϕ***). These samples are obtained by sampling from the prior distribution ***ϕ*** ∼ *p*(***ϕ***) and simulating the model ℳ (***ϕ***). Using the transformation (9), the approximation in (10) can be efficiently evaluated. After training, we performed simulation-based calibration checks to validate the convergence of the neural posterior estimator (Supplementary Figure 1).

### Approximate Bayesian computing with inference-tailored summary statistics

Studies have shown that NPE is sensitive to model misspecification and can potentially provide biased estimates [74, 75]. In contrast, ABC is sensitive to the choice of the summary statistics but converges to the optimal parameter under this summary [76]. To address these limitations, we consider a combined approach in this study: hand-crafted summaries of ABC are replaced by trained summary networks, as in NPE. The learned summaries are computed for each simulation by passing the data through the trained summary network. The *L*^1^ distance is utilized in place of the Wasserstein distance on the learned summaries, as the objective is no longer to compare distributions but rather to enhance robustness against outliers [77]. This approach is referred to as *ABC with inference-tailored summaries* or ABC-NPE.

### Approximate Bayesian computing with posterior mean summary statistics

Previous work suggested training a neural network 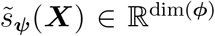 on data ***X*** to approximate the posterior mean 𝔼 [***ϕ*** | ***X***] of the model parameters ***ϕ*** and then using the trained network to compute summary statistics of the data [37, 46]. In this approach, the summary network is trained by optimizing the mean squared error objective

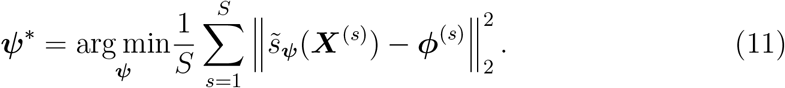

Hence, the output dimension of the network is fixed to the number of parameters. When the acceptance threshold *ϵ* is sufficiently small, this approach, referred to as *ABC with posterior mean summaries* (ABC-PM), converges to a posterior with the same mean as the exact posterior [37, 39].

### Error metrics for posterior and simulated data

We computed the normalized root mean squared error (NRMSE) for each parameter across *N* posterior samples as follows:

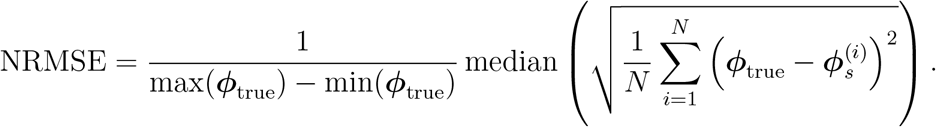

The reported NRMSE is the mean of the parameters.

To quantify the discrepancy between the simulated and observed data, we performed a UMAP projection [78] into a 10-dimensional space and computed the cosine similarity between each simulated and observed cell:

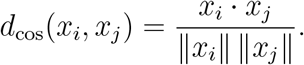

All UMAP hyperparameters were fixed to standard settings.

### Implementation

We used the `FitMultiCell` [79] pipeline to load the model from `Morpheus` [22] and infer the parameters. One simulation takes between 10 s and 30 s. For the ABC algorithm, we chose a sequential Monte Carlo implementation in `pyABC` [80], setting a population size of 1000 particles and using stopping conditions of a minimal acceptance rate of 0.01 or a maximum of 15 generations. The acceptance threshold *ϵ* in each generation was set adaptively by calculating it as the median of the distances from the last population (as it is the default in `pyABC`). The acceptance threshold for the first generation is calculated in a pre-calibration round. We parallelized the simulations in each generation to maximize resource efficiency. If no cells were observed in the simulation, we set the Wasserstein distance to ∞.

We built the neural posterior estimation using `BayesFlow` [81]. Using neural spline flows as an inference network [82], we trained networks with 6 to 8 layers. The smallest network served for inference, while all three together were employed as an ensemble network, i.e., samples of all networks were combined to obtain more conservative posterior estimates, as proposed by [83], and to check for model misspecification [84, 85]. Additionally, we regularized the summary latent space to be normally distributed such that we can check for model misspecification if the real data is far from the simulated data [74].

To extract informative representations from time-resolved cellular data, we designed a summary network composed of convolutional, recurrent, and attention-based components. Each trajectory was first passed independently through a one-dimensional convolutional layer that captured local temporal features. This is followed by a gated recurrent unit (GRU) layer with 32 units, which models sequential dependencies across time points [86]. Both layers operate independently on each cell, ensuring parallel processing across cells. We implemented a temporal attention mechanism to aggregate the latent representations over time. A global query vector was computed by averaging the GRU outputs across all cells. This query is then used to attend to the time steps of each cell, enabling the model to focus on the most informative periods of each trajectory. The attended sequence is subsequently compressed via a pooling operation, resulting in a fixed-dimensional vector (dimension 8) that serves as a population-level summary statistic. Time points were appended to the coordinates, and missing observations in the trajectories (and observations outside the visible window in the experiment) were set to 0 and encoded by an additional binary indicator, as suggested by [87]. This resulted in a 4-dimensional tensor as input to the summary network. The standalone training of the summary network was performed by reducing the summary dimension to the number of parameters and minimizing the mean squared distance of the predictions and parameter values to approximate the posterior mean [37].

We used the same training set for the joint training of the summary and inference networks as for the standalone summary network. We performed 32,000 simulations, taking 1.39 h, employed a batch size of 32, and set 50 as the maximum number of epochs. From a further 100 simulations, we built a validation set and employed an early stopping. All simulations and parameters were standardized by computing the mean and standard deviation of each coordinate and parameter in the validation set. We trained with the Adam optimizer, an initial learning rate of 5 × 10^−4^, and employed a cosine-decay schedule, as it is the default in `BayesFlow`. The training time for the joint training was 2.64 h and for the standalone summary network 4.04 h.

We ran all analyses on a computing cluster using 6 CPU nodes for parallelization, each with 32 cores, and one GPU for training the neural networks. The computing cluster used an Intel Xeon Sapphire Rapids CPU with a core clock speed of up to 2.1 GHz and 60 GB of RAM. The neural network training was performed on a cluster node with an Nvidia A40 graphics card with 48 GB of VRAM.

## Acknowledgments

This work was supported by the German Federal Ministry of Education and Research (BMBF) (EMUNE/031L0293C), the European Union via the ERC grant INTEGRATE, grant agreement number 101126146, and under Germany’s Excellence Strategy by the Deutsche Forschungsgemeinschaft (DFG, German Research Foundation) (EXC 2047—390685813, EXC 2151—390873048, and 524747443), the University of Bonn via the Schlegel Professorship of J.H., and the returning experts fellowship of the Ministry of Innovation, Science, and Research of North-RhineWestphalia (AZ: 421-8.03.03.02-137069). J.M. is a member of the Nanofabrication Facility and is supported by the Institute of Science and Technology Austria. E.K. acknowledges the TRA Life and Health (University of Bonn) as part of the Excellence Strategy of the federal and state governments. We thank Laeschkir Würthner for his insightful comments on the implementation of our model. The views and opinions expressed are those of the authors only and do not necessarily reflect those of the funding agencies.

## Author CRediT

*Conceptualization:* J.A., E.K., J.H.; *Formal analysis:* J.A.; *Investigation:* R.Mo., J.M., E.K.; *Methodology:* J.A., E.A., M.V.; *Resources:* E.K., J.H.; *Software:* J.A., E.A., R.Mu.; *Visualization:* J.A., E.A.; *Writing – original draft:* J.A., E.A.; *Writing – review & editing:* J.A., E.A., R.Mu., M.V., R.Mo., J.M., E.K., J.H..

## Data and code availability

The underlying code and experimental data for this study are available and can be accessed in this repository: github.com/emune-dev/Cell-Migration-Complex-Environments. Additionally, a snapshot of code and data is saved at Zenodo with DOI 10.5281/zenodo.16893454, and the `MorpheusML` model is also provided via this identifier: identifiers.org/morpheus/M6342.

## Competing interests

All authors declare no financial or non-financial competing interests.

## Supporting information

### Additional results for the simulated dataset

**Supplementary Figure 1:**
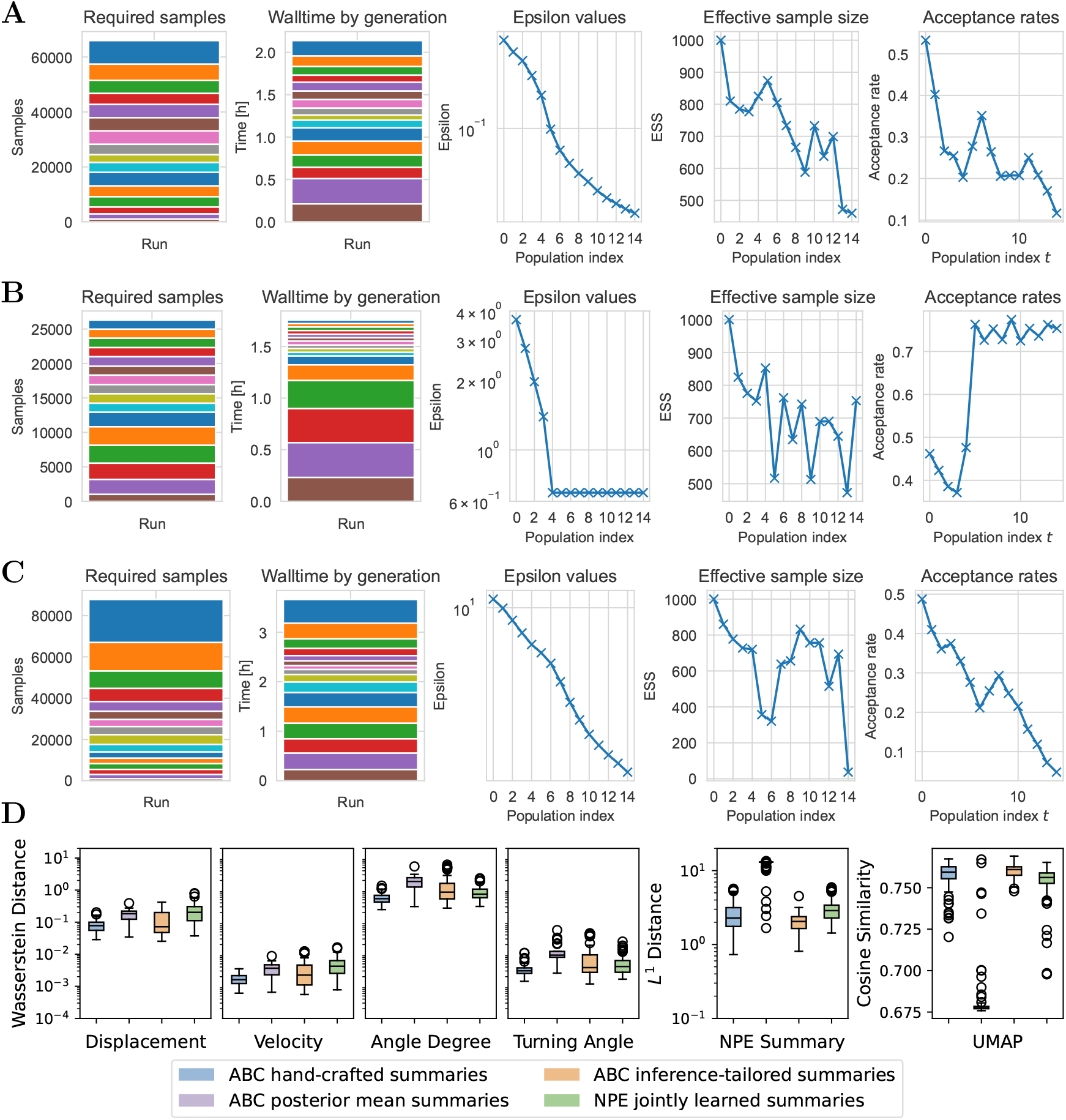
Diagnostics of ABC for each generation of the sampling run on synthetic data set 1. (**A**) Diagnostics of ABC with hand-crafted summaries. (**B**) Diagnostics of ABC-PM with posterior mean summaries. (**C**) Diagnostics of ABC-NPE with inference-tailored summaries. (**D**) Summary statistics for ABC and NPE.

**Supplementary Figure 2:**
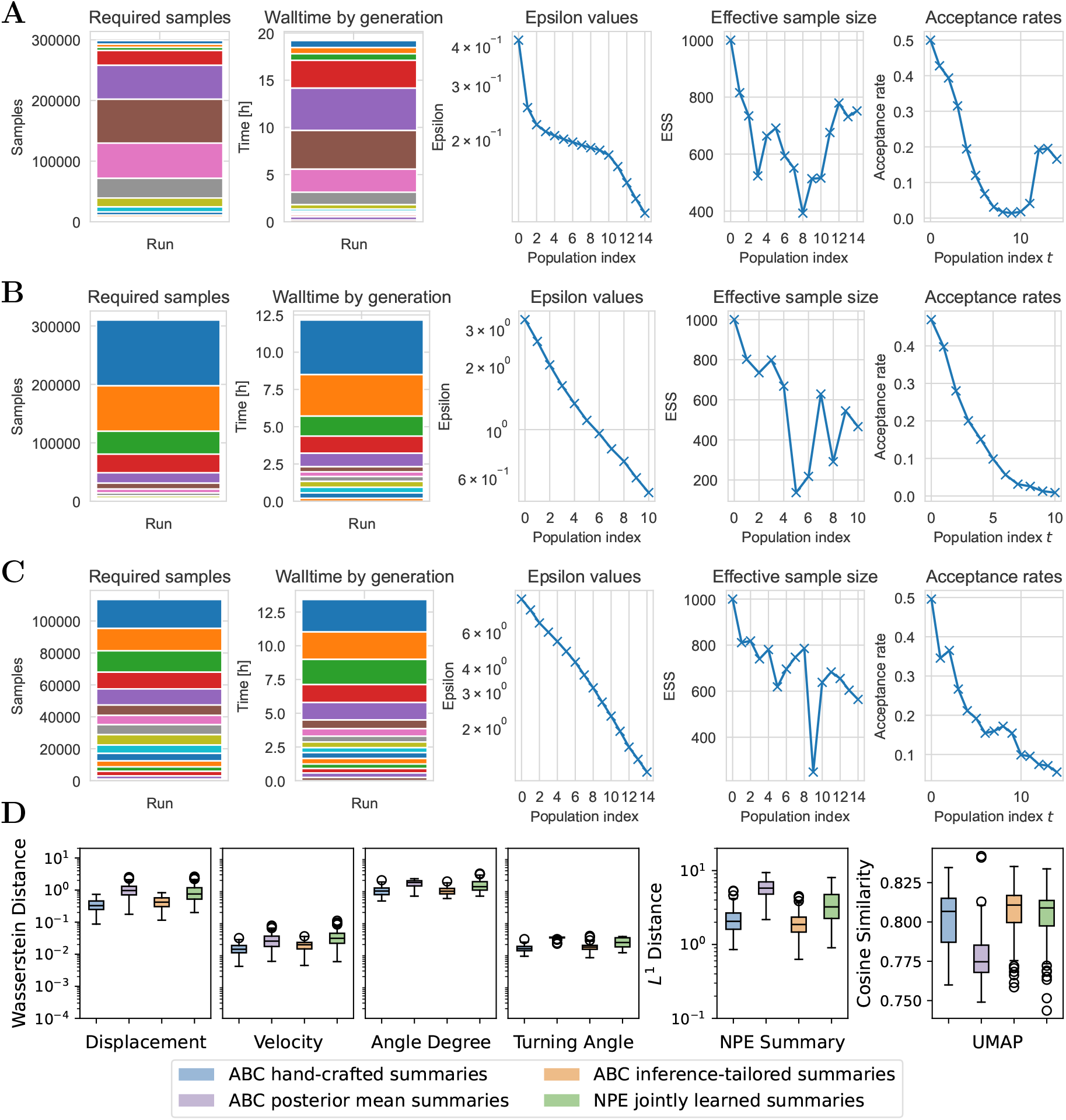
Diagnostics of ABC for each generation of the sampling run on synthetic data set 2. (**A**) Diagnostics of ABC with hand-crafted summaries. (**B**) Diagnostics of ABC-PM with posterior mean summaries. (**C**) Diagnostics of ABC-NPE with inference-tailored summaries. (**D**) Summary statistics for ABC and NPE.

**Supplementary Figure 3:**
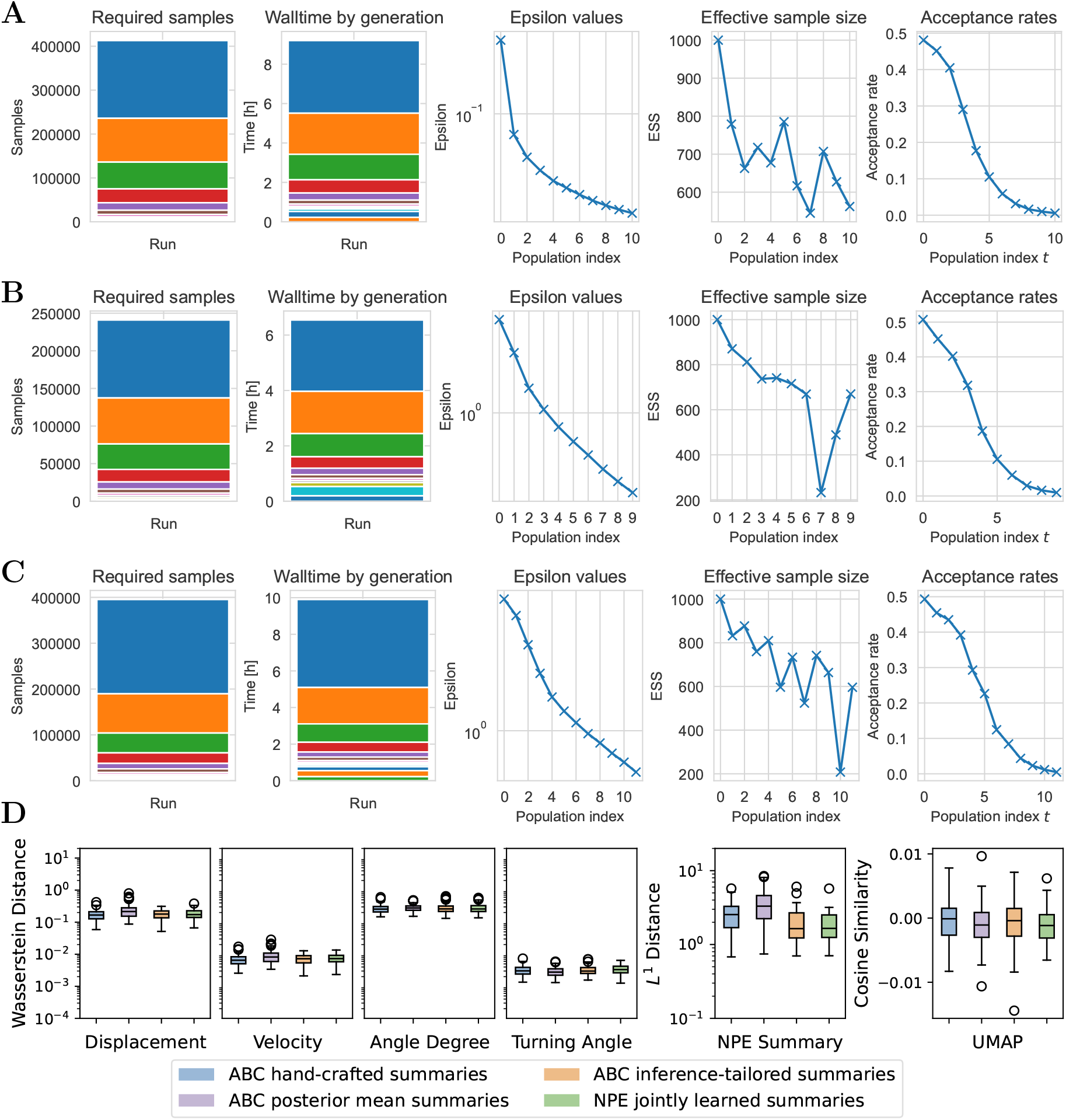
Diagnostics of ABC for each generation of the sampling run on synthetic data set 3. (**A**) Diagnostics of ABC with hand-crafted summaries. (**B**) Diagnostics of ABC-PM with posterior mean summaries. (**C**) Diagnostics of ABC-NPE with inference-tailored summaries. (**D**) Summary statistics for ABC and NPE.

**Supplementary Figure 4:**
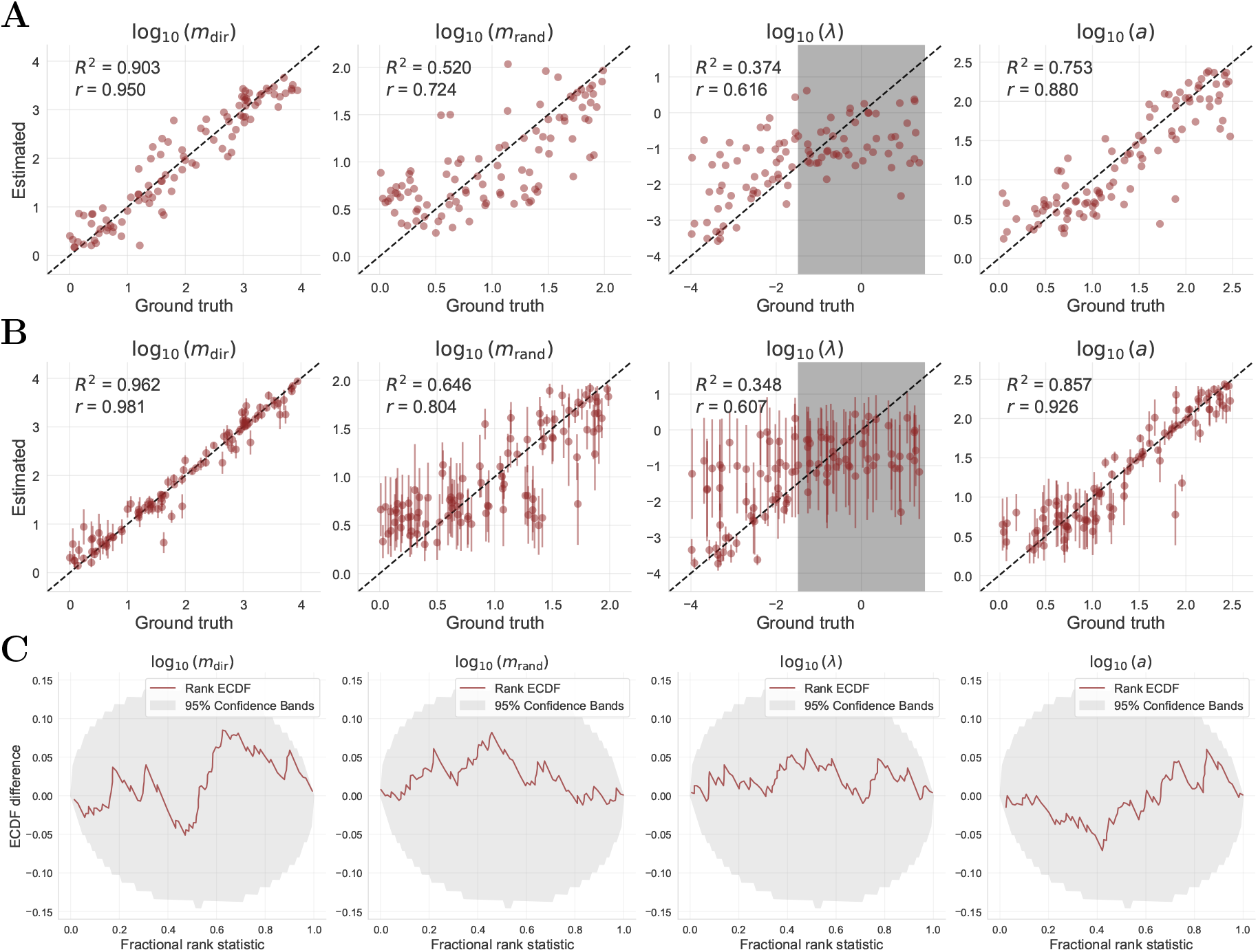
Recovery and empirical cumulative distribution plots. (**A**) Recovery of the parameters by the neural network trained on the posterior mean. The recovery plot displays the posterior median along with the median absolute deviation (if possible). The grey area in the plot of the rate parameter *λ* indicates the area where the mean waiting time 1*/λ* is smaller than the observation time frame of 30 s. (**B**) Recovery of the parameters by the neural posterior estimator. (**C**) The difference between the empirical cumulative distribution (ECDF) of the rank of the true parameter and a uniform ECDF is shown. To assess the validity of the inferred posterior distributions, we employed simulation-based calibration (SBC) [88, 89]. SBC leverages the principle that, under a well-calibrated posterior, the rank of the true parameter within the posterior samples should follow a uniform distribution.

**Supplementary Figure 5:**
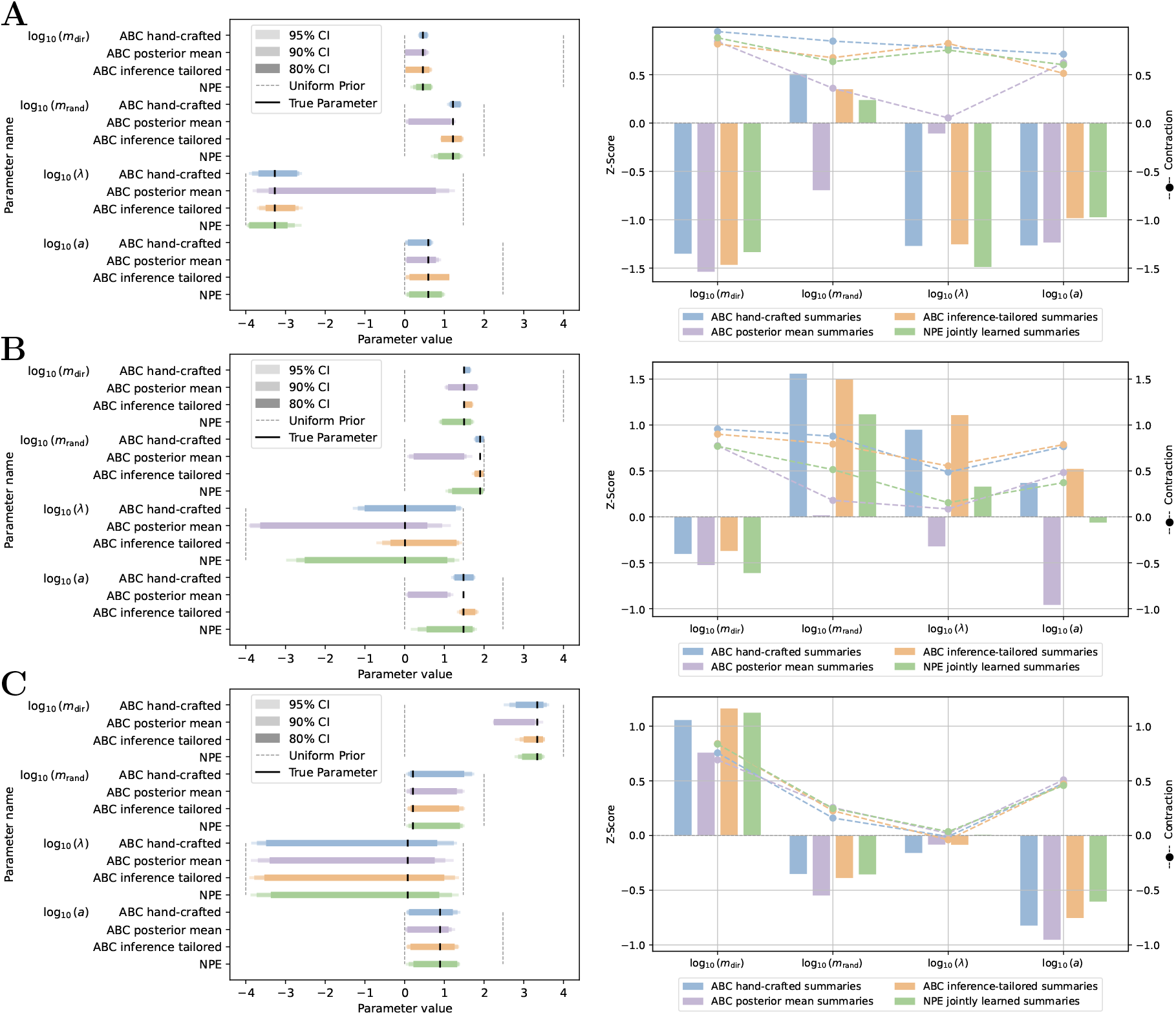
Credible intervals (left) and contraction (right) of the posteriors from the different approaches for the synthetic test data. (**A**) Dataset 1. (**B**) Dataset 2. (**C**) Dataset 3.

**Supplementary Figure 6:**
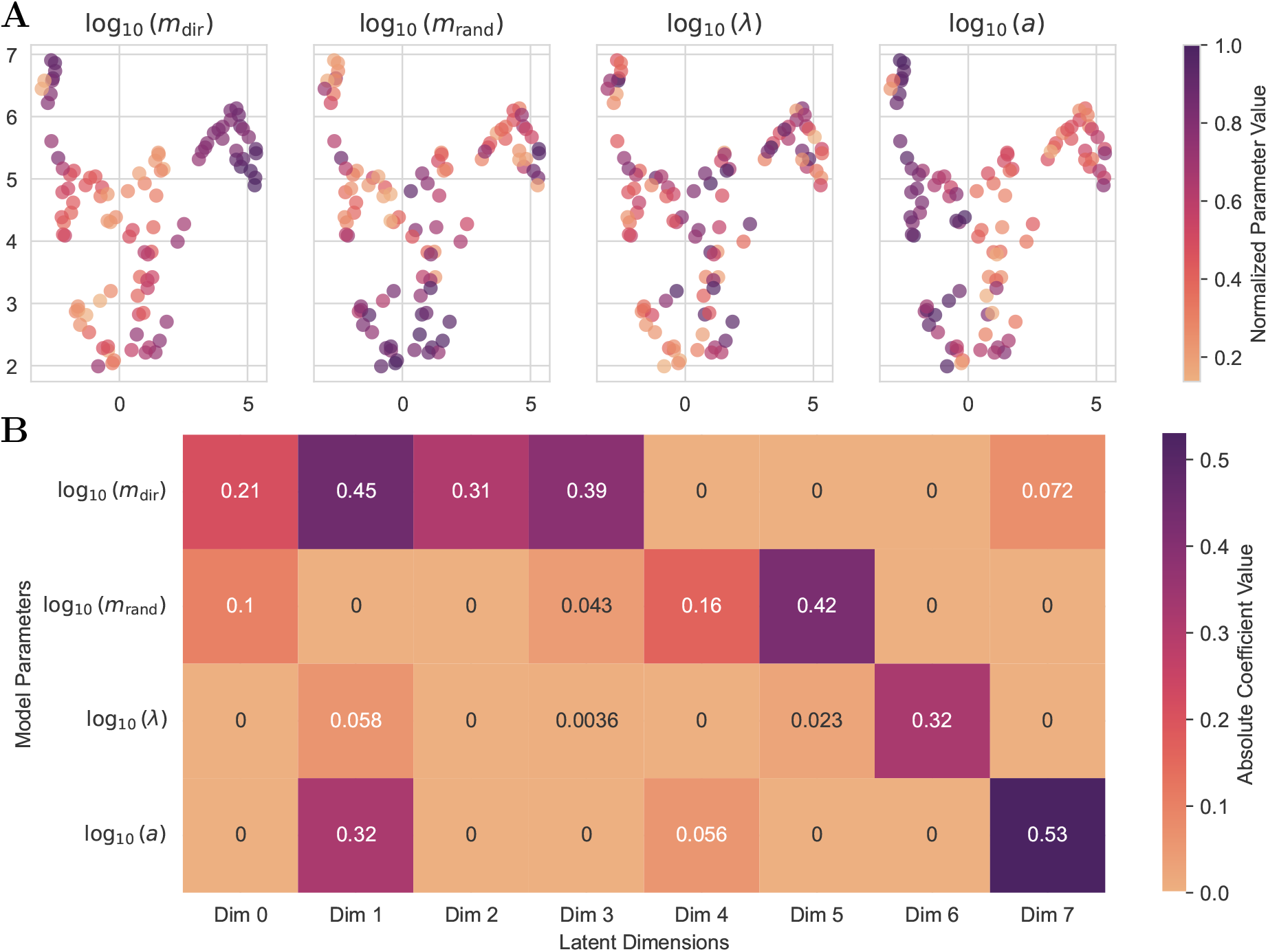
Interpretability of the learned summary space. (**A**) UMAP representation of the learned summary space trained jointly with NPE. (**B**) LASSO-regression (*α* = 0.1) of true parameter values onto the simulation summary space representation reveals that half of the latent dimensions primarily focus on the influence strength of chemokine attraction *m*_dir_, with varying degrees of influence on others.

#### Additional results for the experimental dataset

**Supplementary Figure 7:**
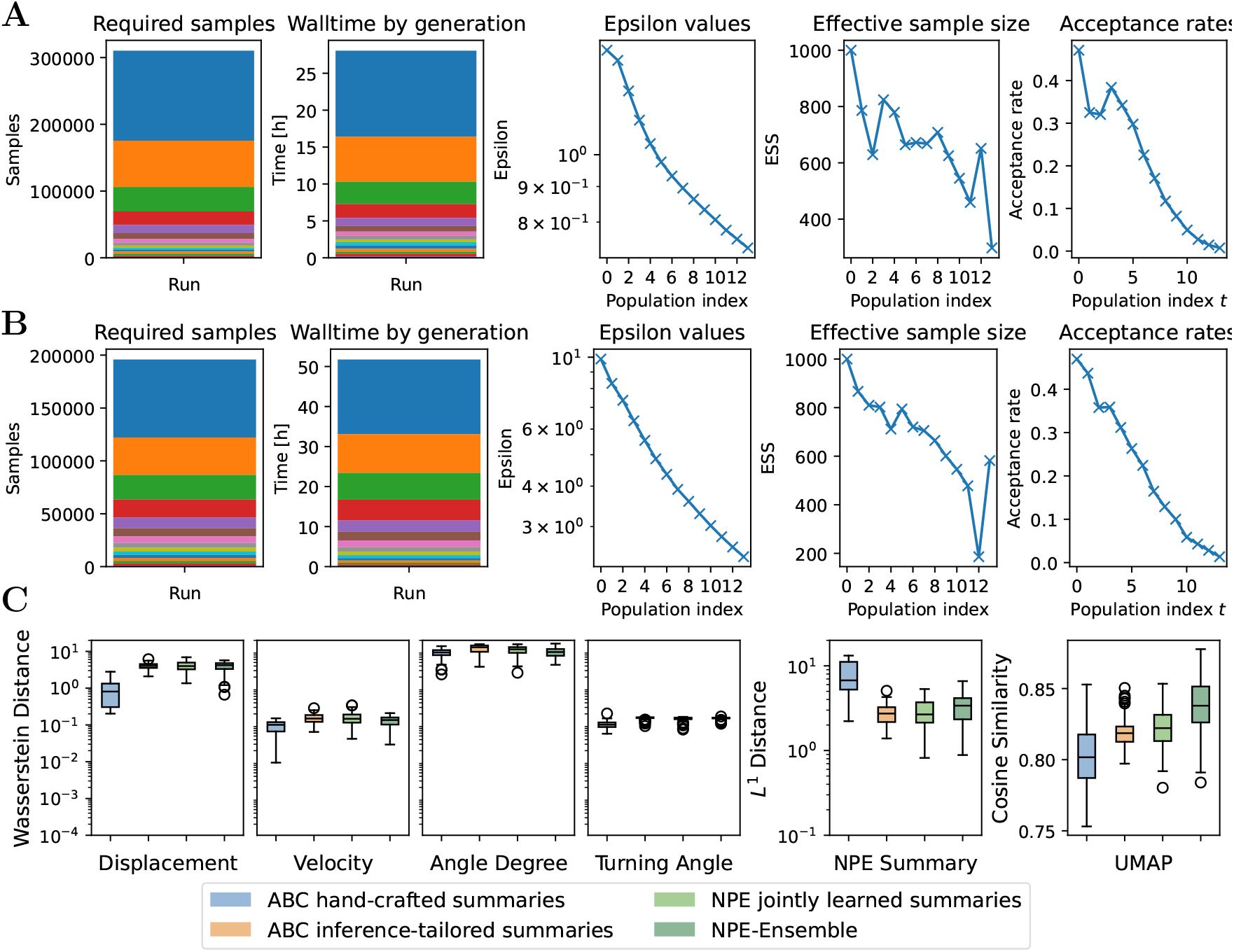
Diagnostics of ABC for each generation of the sampling run on the empirical data. (**A**) Diagnostics for ABC with hand-crafted summaries on experimental data. (**B**) Diagnostics for ABC-NPE with learned summaries on experimental data. (**C**) Summary statistics for ABC and NPE.

Instead of analyzing the full dataset, we can exploit the amortization feature of the neural posterior estimator to estimate the posteriors for truncated datasets, as our summary network accommodates a variable number of cells. We increased the total number of cells in the experiment by randomly subsampling the dataset. Our results were robust to the number of cells used in the experiment (Supplementary Figure 8A).

We can use the learned summary network to compare the summaries of the experimental data with those from the simulations observed during training. During training, we enforced the inlier summary distribution to be Gaussian. Then, misspecification can be detected by a distribution mismatch in the summary space with a high maximum mean discrepancy (MMD), as proposed by [74]. The reference distribution was constructed using the validation data. However, in this case, we cannot reject the hypothesis that the experimental data are consistent with the normal distribution, meaning that no misspecification was detected (see Supplementary Figure 8B).

Additionally, we compute a matrix *K* ∈ ℝ^3×3^ of pairwise Kullback–Leibler divergences between the posteriors inferred by each of the 3 models in the ensemble, as suggested by [85]. Each entry *K*_*ij*_ is estimated by averaging the log-density ratio between models *i* and *j*’s posteriors over samples from model *i*. The magnitude of the entries in *K* serves as a diagnostic tool for ensemble consistency, with small values indicating similar predictive behavior. A heuristic condition for consistency is the normalized maximum divergence max_*i,j*_(*K*_*ij*_*/d*) ≪ 1, where *d* is the dimensionality of the parameter space. In our case, the maximum normalized value is 1.01 for 100 posterior samples, indicating rather high variability between the ensemble members, but no clear evidence of inconsistency within the ensemble.

**Supplementary Figure 8:**
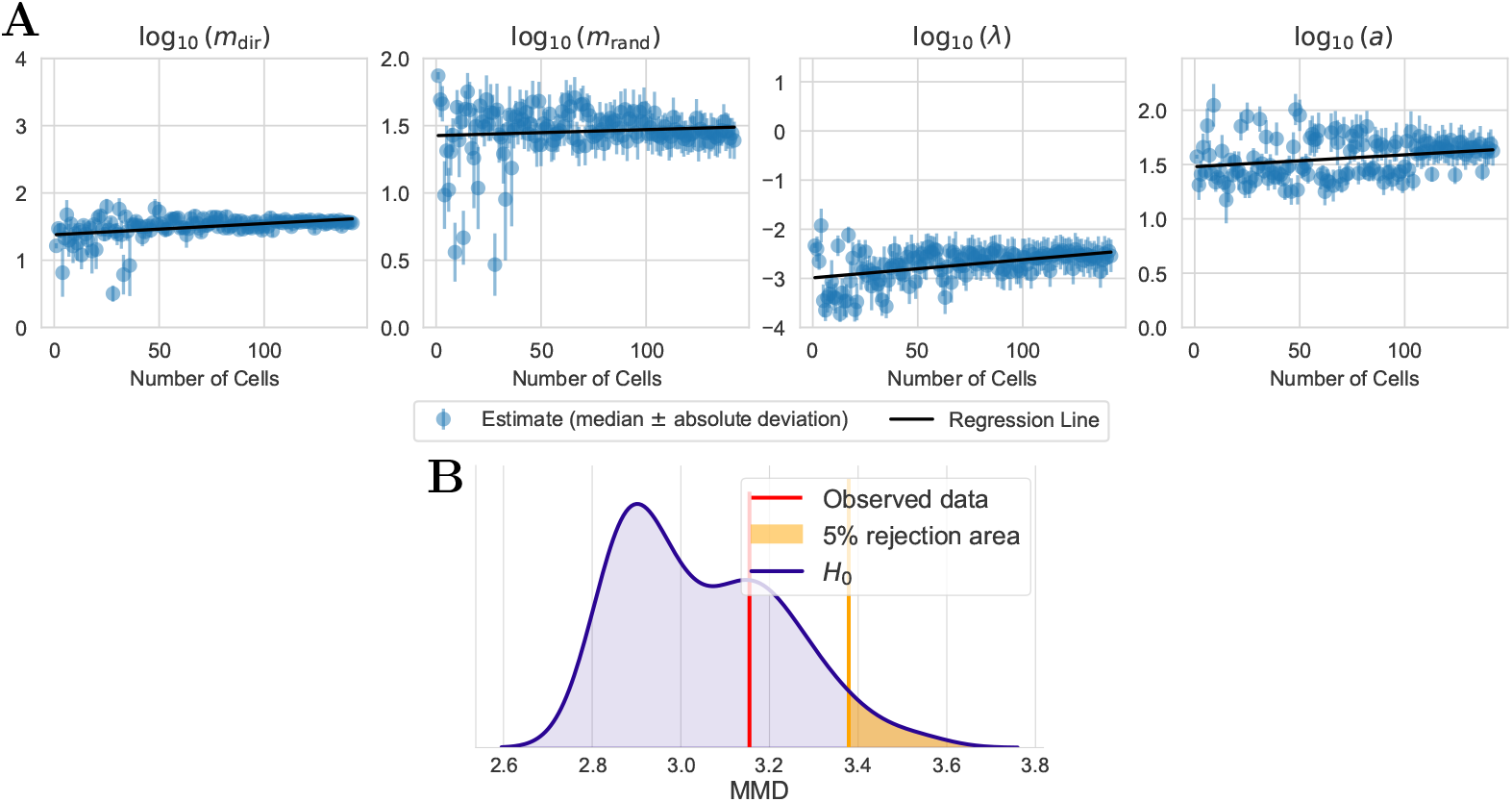
Additional robustness checks. (**A**) Impact of the number of observed cells on the estimated parameters of NPE. (**B**) Model misspecification test using NPE.

#### Additional results for the simulation study

**Supplementary Figure 9:**
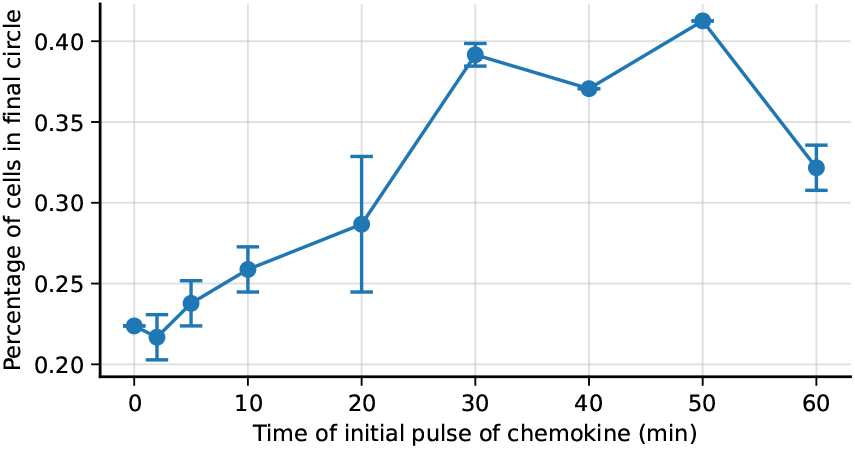
Initial pulse of the chemokine influences trajectory. Median and absolute deviation of the percentage over three simulations from the posterior median.

## Notes

https://github.com/emune-dev/Cell-Migration-Complex-Environments.git

### Competing Interest Statement

The authors have declared no competing interest.

